# Deep Phenotypic Analysis of Blood and Lymphoid T and NK Cells from HIV+ Controllers and Non-Controllers

**DOI:** 10.1101/2021.10.27.466175

**Authors:** Ashley F. George, Xiaoyu Luo, Jason Neidleman, Rebecca Hoh, Poonam Vohra, Reuben Thomas, Min-Gyoung Shin, Madeline J. Lee, Catherine A. Blish, Steven G. Deeks, Warner C. Greene, Sulggi A. Lee, Nadia R. Roan

**Author notes:** **Correspondence:** Sulggi A. Lee. **Correspondence:** Nadia R. Roan.

## Abstract

T and natural killer (NK) cells are effector cells with key roles in anti-HIV immunity, including in lymphoid tissues, the major site of HIV persistence. In this study, we used 42-parameter CyTOF to conduct deep phenotyping of paired blood- and lymph node (LN)-derived T and NK cells from three groups of HIV+ aviremic individuals: elite controllers, and antiretroviral therapy (ART)-suppressed individuals who had started therapy during chronic vs. acute infection, the latter of which is associated with better outcomes. We found that acute-treated individuals are enriched for specific subsets of T and NK cells, including blood-derived CD56-CD16+ NK cells previously associated with HIV control, and LN-derived CD4+ T follicular helper cells with heightened expansion potential. An in-depth comparison of the features of the cells from blood vs. LNs of individuals from our cohort revealed that T cells from blood were more activated than those from LNs. By contrast, LNs were enriched for follicle-homing CXCR5+ CD8+ T cells, which expressed increased levels of inhibitory receptors and markers of survival and proliferation as compared to their CXCR5-counterparts. In addition, a subset of memory-like CD56^bright^TCF1+ NK cells was enriched in LNs relative to blood. These results together suggest unique T and NK cell features in acute-treated individuals, and highlight the importance of examining effector cells not only in blood but also the lymphoid tissue compartment, where the reservoir mostly persists, and where these cells take on distinct phenotypic features.

## 1 Introduction

Lymphocytes with cytotoxic capabilities, including both T and natural killer (NK) cells, are important effectors that play crucial roles in controlling infections by pathogenic viruses, including HIV. T cells include cytotoxic CD8+ T cells, which can directly kill HIV-infected cells, and CD4+ T cells, which can promote antibody responses, enhance CD8+ T cell responses, and directly implement cytotoxic effector functions. NK cells are similarly important in the antiviral immune response against HIV. These classical innate immune cells are rapidly mobilized to recognize and respond to viral infections in a manner regulated by a multitude of activating and inhibitory receptors, and can directly recognize and kill HIV-infected cells (Bjorkstrom et al., 2021).

The activity of T and NK cells against HIV-infected cells is particularly important within lymphoid tissues, where the vast majority of the reservoir persists during antiretroviral therapy (ART) treatment (Estes et al., 2017). Because the follicles of lymphoid tissues appear to harbor a particularly dense array of HIV-infected cells (Brenchley et al., 2012; Lindqvist et al., 2012; Perreau et al., 2013; Banga et al., 2016), strategies are being developed to help direct cytotoxic CD8+ T cell and NK cell populations to this site (Carrega and Ferlazzo, 2012; Leong et al., 2016). These strategies include creating HIV/SIV-specific CAR (chimeric antigen receptor) T cells engineered to express CXCR5, a chemokine receptor mediating entry into the follicles (Ayala et al., 2017; Haran et al., 2018), or IL-15:IL-15Rα-based therapies, which upregulate CXCR5 in cytotoxic CD8+ T cells, thus facilitating their access to follicles (Watson et al., 2018; Webb et al., 2018). People living with HIV (PLWH) harbor HIV-specific CXCR5+ CD8+ T cells both in circulation as well as within lymphoid tissues. These follicular HIV-specific T cells appear to have poor cytolytic activity compared to their blood counterparts (Reuter et al., 2017; Nguyen et al., 2019). Nonetheless, follicular CXCR5+ CD8+ T cells from LNs may contribute to viral control, since the viral load in PLWH correlates negatively with the frequency of these cells (Reuter et al., 2017; Nguyen et al., 2019). However, beyond the observation that these cells are poorly cytotoxic within the LNs, other features of these cells are poorly understood.

NK cells, like CXCR5+ CD8+ T cells, may also be important in controlling HIV within B cell follicles. In SIV-infected African green monkeys that are able to control the infection without any treatment, lymphoid NK cells were largely CXCR5+ and persisted within the follicles of LNs (Huot et al., 2017). When NK cells were depleted in these animals via anti-IL-15 antibody, viremia increased in lymph nodes, suggesting an important role for follicular NK cells in viral control. In addition to classical innate responses, NK cells can also mount “memory” recall responses, including against HIV (Peppa et al., 2018; Nikzad et al., 2019; Vendrame et al., 2020; Wang et al., 2020). One subset of memory-like NK cells, significantly increased in the peripheral blood of PLWH, was found to develop and expand in response to inflammatory cytokines, including IL-15 (Wang et al., 2020). This expansion was mediated by *TCF7* (transcript for TCF1), a transcriptional factor important for cellular quiescence and stemness (Kuo and Leiden, 1999; Zhou et al., 2010). Identified as CD94+*TCF7*+CD56^bright^ NK cells, these memory-like NK cells exhibit increased rates of proliferation and more robust effector functions in response to stimulation, and may aid in HIV control.

Unfortunately, both T and NK cells’ effector responses often become dysfunctional upon HIV infection (Hu et al., 1995; Betts et al., 2006), and do not completely recover even after ART treatment (Korencak et al., 2019; Nabatanzi et al., 2019), which historically has been initiated during the chronic phase of infection. However, there exists rare PLWH that can naturally control HIV in the absence of ART through immune-mediated mechanisms. These include elite controllers, who are thought to control the virus primarily through expression of specific class-I HLA alleles, which are able to elicit protective anti-HIV CD8+ T cell responses (Migueles et al., 2008; International et al., 2010; Avettand-Fenoel et al., 2019; Gaiha et al., 2019; Monel et al., 2019; Casado et al., 2020). More recently, post-treatment controllers (PTCs) have been described that exhibit durable control of HIV following withdrawal of ART. These individuals don’t appear to express the protective class-I HLA alleles found in elite controllers (Saez-Cirion et al., 2013), and may instead use NK cell-mediated mechanisms (Scott-Algara et al., 2015). Interestingly, PTCs are over-represented among PLWH who initiate ART during acute infection (Lodi et al., 2012; Saez-Cirion et al., 2013; Maenza et al., 2015; Sneller et al., 2017). In addition, PLWH who initiate ART during acute infection have lower levels of T cell activation (Jain et al., 2013) and a smaller HIV reservoir that decays more rapidly (Chun et al., 2007; Hocqueloux et al., 2013), relative to those who initiated treatment during chronic infection. This is likely due to better preserved immune responses in these individuals (Takata et al., 2017; Ndhlovu et al., 2019; Kazer et al., 2020; Leyre et al., 2020). While early ART administration is now standard in the field, relatively little is known about the features of effector cells in these individuals and to what extent they may differ from those of PLWH who initiated treatment during chronic infection. A better understanding of the features of effector cells from PLWH who initiated ART during acute infection may provide insights into the basis of a better-preserved anti-HIV immune response and post-treatment control of HIV.

Overall, these observations suggest different ways in which HIV-infected people achieve full or partial immunological control of their infection. Understanding the basis of their success could inform strategies to eliminate infected cells or improve PLWH’s ability to control HIV after ART cessation. As a step towards these goals, we undertook a deep assessment of effector T and NK cells from three categories of aviremic PLWH: elite controllers, and virally-suppressed non-controllers who had started therapy either during the acute or the chronic stage of infection. We designed a 42-parameter CyTOF panel, tailored for simultaneous characterization of T and NK cells, and implemented it on freshly isolated paired blood and lymphoid tissue specimens from PLWH. By comparing across participant groups, and between sampling sites, we define characteristics of T and NK cells that associate with a specific state of HIV control, and identify profound phenotypic differences between effector cells from blood and lymphoid tissue.

## Materials and Methods

### 1.1 Study Participants and Specimen Collection

The study was approved by the University of California, San Francisco (UCSF) under IRB # 10– 01330, and all participants provided informed consent. Blood was drawn from 19 participants total, and for 13 participants, LN specimens were additionally collected under local anesthesia using ultrasound-guided fine need aspiration (FNA) at the same study visit. No adverse effects from FNA procedures were reported, and participants reported minimal to no discomfort during and after the procedure. Demographical and clinical characteristics of the study’s HIV+ participants are summarized in Supplementary Table S1. Participants included: 1) 5 elite controllers, 2) 8 individuals on long-term ART who initiated therapy during acute (<1 year) infection, and 3) 6 individuals on long-term ART who initiated therapy during chronic (>1 year) infection (Supplementary Table S1). All 14 individuals on ART were stably suppressed: their HIV-1 RNA load was <110 copies/ml (<40 HIV-1 RNA copies/ml in 12 participants) and their CD4 T cell count >400 cells/μL.

### 1.2 Isolation of Cells

PBMCs were isolated from whole-blood specimens using Lymphoprep density gradient medium according to the manufacturer’s protocol (StemCell Technologies, Vancouver, BC, Canada). LN specimens were obtained by FNA biopsies of 1–2 lymph nodes in the inguinal chain area. Generally, 4–5 passes per LN were performed using 3 cc slip tip syringes and 23- or 22-gauge needles of 1 ½ inch in length. To confirm LN sample acquisition, a portion of the first pass was fixed in 95% alcohol on a slide, stained with Toluidine Blue, and examined by microscopy. Following confirmation from the cytopathologist, the remaining sample was transferred into RPMI medium (Corning, Corning, NY, USA). LN specimens were mechanically dissociated into single-cell suspension using a 5 ml syringe and a 70-micron strainer.

### 1.3 Preparation of Cells for CyTOF

Freshly isolated cells from blood and LN specimens were immediately treated with cisplatin (Sigma-Aldrich, St. Louis, MO, USA) as a live/dead marker and then fixed with paraformaldehyde as described previously (Cavrois et al., 2017; Ma et al., 2020; Neidleman et al., 2020a; Neidleman et al., 2020b; Neidleman et al., 2021; Xie et al., 2021). Briefly, a total of 6 × 10^6^ cells were resuspended in 2 ml PBS (Rockland, Gilbertsville, PA, USA) with 2 mM EDTA (Corning, Corning, NY, USA). Then, 2 ml of PBS with 2 mM EDTA and 25 μM cisplatin (Sigma-Aldrich, St. Louis, MO, USA) was added to the cells, quickly mixed, and the mix incubated for 60 s at room temperature. Cisplatin staining was quenched with 10 ml CyFACS (metal contaminant-free PBS [Rockland, Gilbertsville, PA, USA] supplemented with 0.1% bovine serum albumin [BSA; Sigma-Aldrich, MO, USA] and 0.1% sodium azide [Sigma-Aldrich, St. Louis, MO, USA]). Following centrifugation, cells were resuspended in 2% paraformaldehyde (PFA, Electron Microscopy Sciences, Hatfield, PA, USA) in CyFACS and incubated for 10 min at room temperature. Cells were washed twice in CyFACS, resuspended in 100 μL of CyFACS containing 10% DMSO, and stored at −80°C until analysis by CyTOF.

### 1.4 CyTOF Staining and Data Acquisition

Staining of cells for CyTOF analysis was conducted as previously described (Cavrois et al., 2017; Ma et al., 2020; Neidleman et al., 2020a; Neidleman et al., 2020b; Neidleman et al., 2021; Xie et al., 2021). To minimize cell loss, multiple cisplatin-treated samples were barcoded and combined using the Cell-ID 20-Plex Pd Barcoding Kit (Fluidigm, South San Francisco, CA, USA), according to the manufacturer’s protocol. Briefly, the fixed samples were thawed, washed twice with Maxpar Barcode Perm Buffer (Fluidigm, South San Francisco, CA, USA), and resuspended in 800 μL of Maxpar Barcode Perm Buffer in Nunc 96 DeepWell polystyrene plates (Thermo Fisher, Waltham, MA, USA). Each barcode (10 μL) was diluted in 100 μL of Maxpar Barcode Perm Buffer, added to each sample, and the mix incubated for 30 min at room temperature. Individual samples were washed once with MaxPar Cell Staining Buffer (Fluidigm, South San Francisco, CA, USA), once with CyFACS, and then combined.

Barcoded cells were aliquoted at a concentration of 6 × 10^6^ cells/800 μL CyFACS per well into Nunc 96 DeepWell polystyrene plates. Cells were then blocked with 1.5% mouse sera (Thermo Fisher, Waltham, MA, USA), 1.5% rat sera (Thermo Fisher, Waltham, MA, USA), and 0.3% human AB sera (Sigma-Aldrich, St. Louis, MO, USA) for 15 min at 4°C. Cells were then washed twice with CyFACS and incubated for 45 min at 4°C in 100 μL/well with the cocktail of surface antibodies (Supplementary Table S2). Cells were then washed 3X with CyFACS and incubated at 4°C overnight in 2% PFA (Electron Microscopy Services, Hatfield, PA, USA) in PBS (Rockland, Gilbertsville, PA, USA). Cells were then incubated with Intracellular Fixation and Permeabilization Buffer (eBioscience, San Diego, CA, USA) for 30 min at 4°C. Cells were then washed twice in Permeabilization Buffer (eBioscience, San Diego, USA) and blocked for 15 min at 4°C in 100 μL/well of 0.15% mouse and 0.15% rat sera diluted in Permeabilization Buffer. After washing once with Permeabilization Buffer, cells were incubated for 45 min at 4°C in 100 μl/well with the cocktail of intracellular antibodies (Supplementary Table S2) diluted in Permeabilization Buffer. Cells were then wash once with CyFACS and incubated for 20 min at room temperature with 250 nM Cell-ID Intercalator-IR (Fluidigm, South San Francisco, CA, USA). Afterwards, cells were washed twice with CyFACS and incubated overnight at 4°C in CyFACS. Prior to sample acquisition, cells were washed once with MaxPar Cell Staining Buffer (Fluidigm, South San Francisco, CA, USA), once with Cell Acquisition Solution (Fluidigm, South San Francisco, CA, USA), and then resuspend in 1X EQ Four Element Calibration Beads (Fluidigm, South San Francisco, CA, USA) diluted in Cell Acquisition Solution (Fluidigm, South San Francisco, CA, USA). The concentration of cells was adjusted to acquire at a rate of ∼300 events/sec using a wide-bore (WB) injector on a Helios-upgraded CyTOF2 instrument (Fluidigm, South San Francisco, CA, USA) at the UCSF Parnassus Flow Core Facility.

### 1.5 CyTOF Data Analysis

CyTOF datasets were normalized and de-barcoded according to the manufacturer’s protocol (Fluidigm, South San Francisco, CA, USA). FlowJo software (BD Biosciences) was used to gate for events corresponding to total T cells (live, singlet, CD19-CD14-CD33-CD3+), CD4+ T cells (live, singlet, CD19-CD14-CD33-CD3+CD8-), CD8+ T cells (live, singlet, CD19-CD14-CD33-CD3+CD8+), and NK cells (CD19-CD14-CD33-CD3-CD4-CD7+) among PBMC and LN specimens (Fig. S1). For data visualization, tSNE analyses were performed in Cytobank software (Beckman Coulter, Brea, CA, USA) with default settings. Markers used in the upstream gating strategy and non-cellular markers (e.g., live/dead stain, Pd barcodes) were excluded as tSNE parameters. Dot plots were created using FlowJo (Ashland, OR, USA). The statistical tests used in comparison of groups are indicated within the figure legends.

For tSNE clustering of datasets comparing effector cells from the three patient groups, the Seurat R package for single-cell analysis (Satija et al., 2015) was used to define clusters of cells based on 39 CyTOF parameters, excluding CD19, CD14, and CD33, and clusters were filtered based on a minimum cell-count threshold of 50. For blood-derived T cells, 40,000 cells were down-sampled (randomly selected) to enable processing within memory limits and computational efficiency. A total of 11,338 LN NK cells, 240,804 blood NK cells, 463,722 LN T cells, and 759,966 blood T cells, were subjected to Seurat clustering. CyTOF clusters were identified using a shared nearest neighbor (SNN) graph based on the Jaccard index (Levine et al., 2015). The optimal clustering resolution parameters were determined using Random Forest (Shin et al., 2019) and a silhouette score-based assessment of clustering validity and subject-wise cross-validation. The silhouette score is a metric that assesses cluster validity based on within cluster distance and between cluster distance, and is often used to find the best number of clusters (Lovmar et al., 2005). The efficacy of resolution parameters ranging from 0.01 to 0.2 were tested, and silhouette scores were measured from each case. Cross-validation was used to assure the reproducibility of the optimization process. For the cross-validation, CyTOF profiles were used as input features and cluster memberships were used as labels to train a Random Forest model using N-1 subjects, where N is the total number of subjects. The rest of the data were used to predict the cells’ cluster level and calculate the silhouette scores. This cross-validation procedure was performed N times, iterating through each of the subjects. Silhouette score was first averaged across cells in the same cluster and then the mean of the averaged scores was used as the final score. The optimal resolution parameter was selected based on the silhouette score trend across different resolution parameter values, using the elbow method. Finally, a generalized linear mixed model (GLMM) was used to estimate the association between cluster membership and infection status. The log odds ratio was retrieved from the coefficient estimate of each variable. To assess the association between CyTOF marker expression quantile levels and group (controller, acute-treated, chronic-treated), a linear model analysis was performed. All p-values were corrected using the FDR, and a threshold of 0.05 was used to determine significance. GLMM was performed using R package lme4 (Bates et al., 2015), and linear regression was performed using the lm function in R.

### 1.6 Data Availability Statement

The raw datasets generated for this study can be found in the public repository Dryad, and accessible at the following link: https://doi.org/10.7272/Q6CV4FZJ.

## 2 Results

### 2.1 Patient population and specimens

To characterize T and NK cells from aviremic PLWH, we recruited 13 HIV+ participants who donated both PBMCs and LN aspirates during the same study visit, along with an additional 6 participants who donated PBMCs alone (Supplementary Table S1). Participants consisted of: elite controllers who were not taking ART (n=5), participants who initiated ART during acute HIV (<150 days from estimated date of infection) and were suppressed for at least 12 months at the time of sampling (acutely-treated, n=8), and participants who initiated ART during chronic HIV (>2 years from infection) and were suppressed for at least 12 months at the time of sampling (chronically-treated, n=6). Eligibility criteria for all participants were: (1) confirmed HIV-1 infection and (2) undetectable HIV-1 (<40 copies RNA/ml) for at least 12 months. For HIV non-controllers, an additional eligibility criterion was 3) at least 12 months of continuous ART at study entry with no regimen changes in the preceding 24 weeks. Excluded participants were those with (1) recent hospitalization, (2) recent infection requiring systemic antibiotics, (3) recent vaccination or (4) exposure to any immunomodulatory drug (including maraviroc) in the 16 weeks prior to study. Participants were consented by the investigator before any procedures took place, eligibility was confirmed, and medical history and concurrent medications ascertained. The research was approved by the University of California Committee on Human Research (CHR), and all participants provided written informed consent.

Cells were processed immediately after collection, stained with cisplatin as a viability marker, fixed with paraformaldehyde (PFA), and frozen until all donor samples were acquired. A 42-parameter CyTOF panel was established to simultaneously phenotype T and NK cells for markers of differentiation and activation states, homing receptors, and inhibitory and activating NK cell receptors (Supplementary Table S2). Importantly, this panel enabled analysis of PFA-fixed cells that never went through cryopreservation. To avoid batch effects, all samples were barcoded, stained, and run on the mass cytometer together (Materials and Methods).

### 2.2 Elite Controllers and Acutely-treated PLWH Have More Naïve T Cells in Peripheral Blood than Chronically-treated PLWH

We first conducted a systematic comparison of lymphocytes between the participant groups. We applied t-distributed stochastic neighbor embedding (tSNE) to qualitatively assess for overall phenotypic similarities and differences between CD4+ T cells, CD8+ T cells, and NK cells from PBMCs (Fig. 1A) and lymph nodes (LNs; Fig. 1B) between the 3 patient groups. Although CD4+ T cells, CD8+ T cells, and NK cells resided in distinct areas of the tSNE, these subsets occupied similar areas in the 3 participant groups. However, there were subtle differences in tSNE distribution of the T cells. In blood, there were two regions of the tSNE (Fig. 1A, circled in black) occupied by CD8+ T cells from controllers, one of which was almost entirely absent in acutely-treated individuals, and both of which were relatively devoid in chronically-treated individuals. In LNs, one distinct region occupied by CD4+ T cells (Fig. 1B, circled in black) in the controllers and acutely-treated individuals, was devoid of cells in chronically-treated individuals. These results suggest some phenotypic differences in T cells between controllers, acutely-treated individuals, and chronically-treated individuals in both blood and LNs.

**Figure 1.**
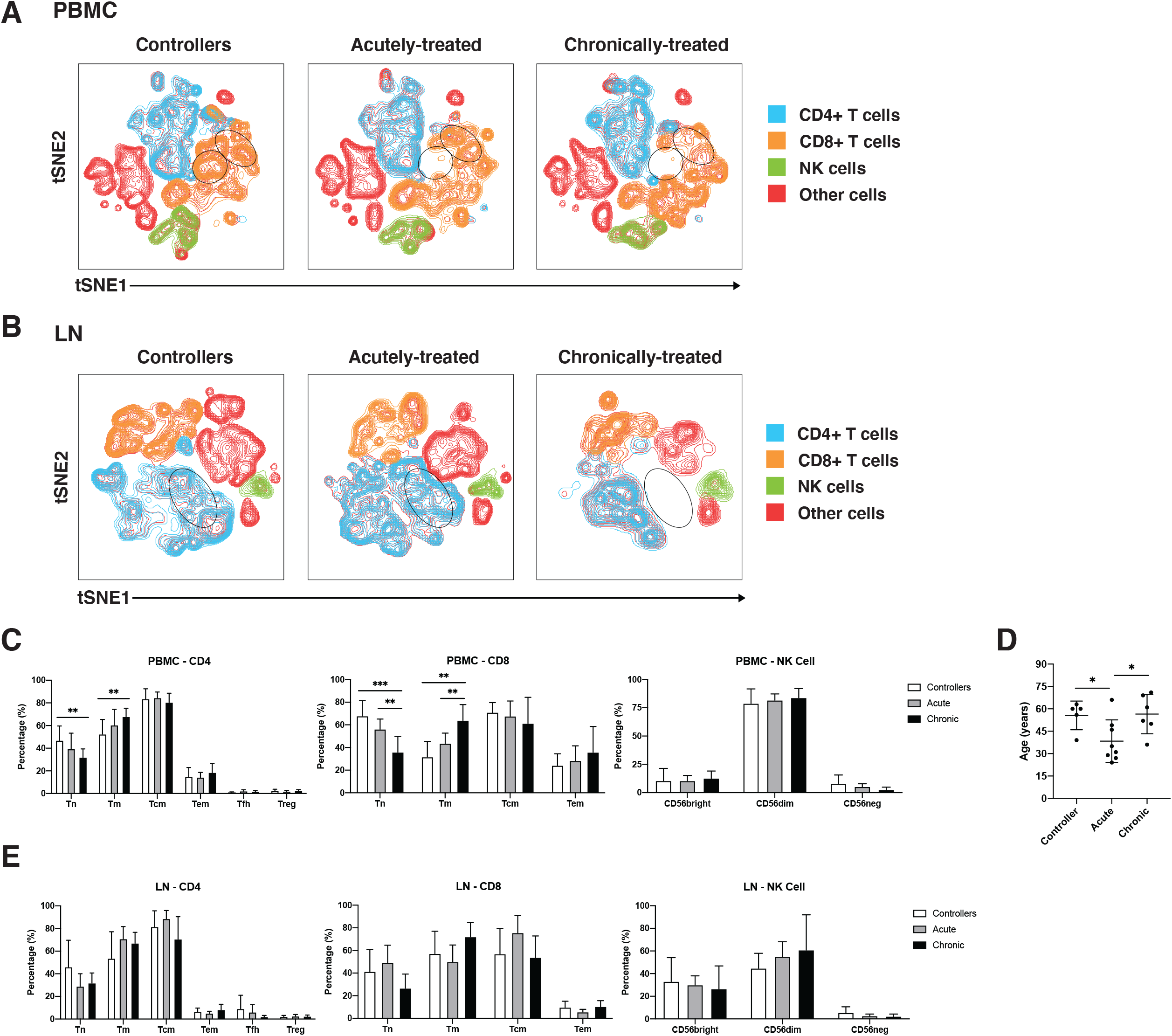
Characterization of T and NK cells from PBMCs and LNs of HIV controllers and non-controller HIV+ individuals treated during acute or chronic infection. **(A, B)** t-distributed stochastic neighbor embedding (tSNE) plots for T and NK cells from PBMCs (A) and LNs (B) of HIV controllers (left), acutely-treated (middle), and chronically-treated (right) HIV+ individuals. The main subsets analyzed are each colored as indicated. Some subsets of T cells were preferentially detected in the controllers (regions circled in black). **(C)** Blood naïve T cells are more frequent in controllers than non-controllers. Shown are frequencies of canonical subsets in PBMCs from the indicated groups of HIV+ individuals among CD4+ T cells (live, singlet, CD19-CD14-CD33-CD3+CD8-, left), CD8+ T cells (live, singlet, CD19-CD14-CD33-CD3+CD8+, middle) and NK cells (CD19-CD14-CD33-CD3-CD4-CD7+, right). Subsets of CD4+ and CD8+ T cells were defined as follows: naïve T cells (Tn): CD45RO-CD45RA+; memory T cells (Tm): CD45RO+CD45RA-; central memory T cells (Tcm): CD45RO+CD45RA-CCR7+CD27+; and effector memory T cells (Tem): CD45RO+CD45RA-CCR7-CD27-. In addition, CD4+ T cells were further subsetted into T follicular helper cells (Tfh): CD45RO+CD45RA-CXCR5+PD1+; and regulatory T cells (Treg): CD45RO+CD45RA-CD25+CD127-. Subsets of NK cells were defined based on CD56^bright^, CD56^dim^, and CD56-staining. **(D)** Age distribution of participants in each group. Controllers were not younger than non-controllers. **(E)** Frequencies of canonical T and NK cell subsets from LNs are similar between controllers and non-controllers. Frequencies of the indicated subsets of CD4+ T cells (left), CD8+ T cells (middle), and NK cells (right) in LNs from the indicated groups are shown. *p<0.05, **p<0.01, ***p<0.001 as assessed using the Student’s unpaired t test.

To investigate these differences in a more quantitative manner, we assessed the composition of the CD4+ T cell (naïve, Tn; memory, Tm; central memory, Tcm; effector memory, Tem; T follicular helper cells, Tfh; regulatory T cells, Treg), CD8+ T cell (Tn, Tm, Tcm, Tem), and NK cell subsets from PBMCs (Fig. 1C) and LNs (Fig. 1G), with the three main subsets of NK cells being classically defined by CD56 expression (CD56^bright^, CD56^dim^, CD56-) (Vendrame et al., 2020; Zhao et al., 2020). CD4+ and CD8+ Tn cells were significantly more frequent in the PBMCs of controllers than of chronically-treated individuals (Fig. 1C). CD8+ Tn cells in PBMCs were also significantly more frequent in the acutely-treated than the chronically-treated group. Conversely, blood CD4+ and CD8+ Tm cells were more frequent in the chronically-treated group relative to the other two groups. As the proportion of Tn to Tm cells decreases with age (Cossarizza et al., 1996; Saule et al., 2006), we assessed whether the increase in Tn proportions in our controllers were accounted by younger age. In fact, our controllers were on average older than the acutely-treated participants (Fig. 1D), suggesting that age did not account for our observations. In contrast to data from the blood, the frequencies of the analyzed CD4+, CD8+, or NK cell subsets from the LNs did not show any significant differences between the patient groups in LNs (Fig. 1E). Together, these results indicate that in blood, controllers and acutely-treated individuals have an increased frequency of Tn cells compared to chronically-treated individuals on ART.

### 2.3 High-dimensional Clustering Reveals Unique T and NK Cell Phenotypes Associated with Acutely-treated PLWH

To take advantage of all phenotyping markers from our high-dimensional CyTOF datasets, we defined T and NK cell subsets via unsupervised clustering. Four clusters each of T cells and NK cells were identified from PBMCs, and six T cell and four NK cell clusters from LNs (Fig. 2A). We then used a generalized linear mixed model (GLMM) to estimate the association between cluster membership and group. Clusters significantly associated with a specific state of HIV control were identified for blood T cells, blood NK cells, and LN T cells (Fig. 2A). In particular, T cell Cluster 3 from blood (Acute vs. Controllers: P < 0.0001; Acute vs. Chronic: P < 0.009) and T cell Cluster 9 from LNs (Acute vs. Controllers: P < 0.03; Acute vs. Chronic: P < 0.0001) were enriched in acutely-treated individuals compared to controllers and the chronically-treated individuals. In addition, NK cell Cluster 8 from blood was enriched in acutely-treated individuals as compared to chronically-treated ones (P < 0.05). No significant associations with disease state were found for LN NK cells.

**Figure 2.**
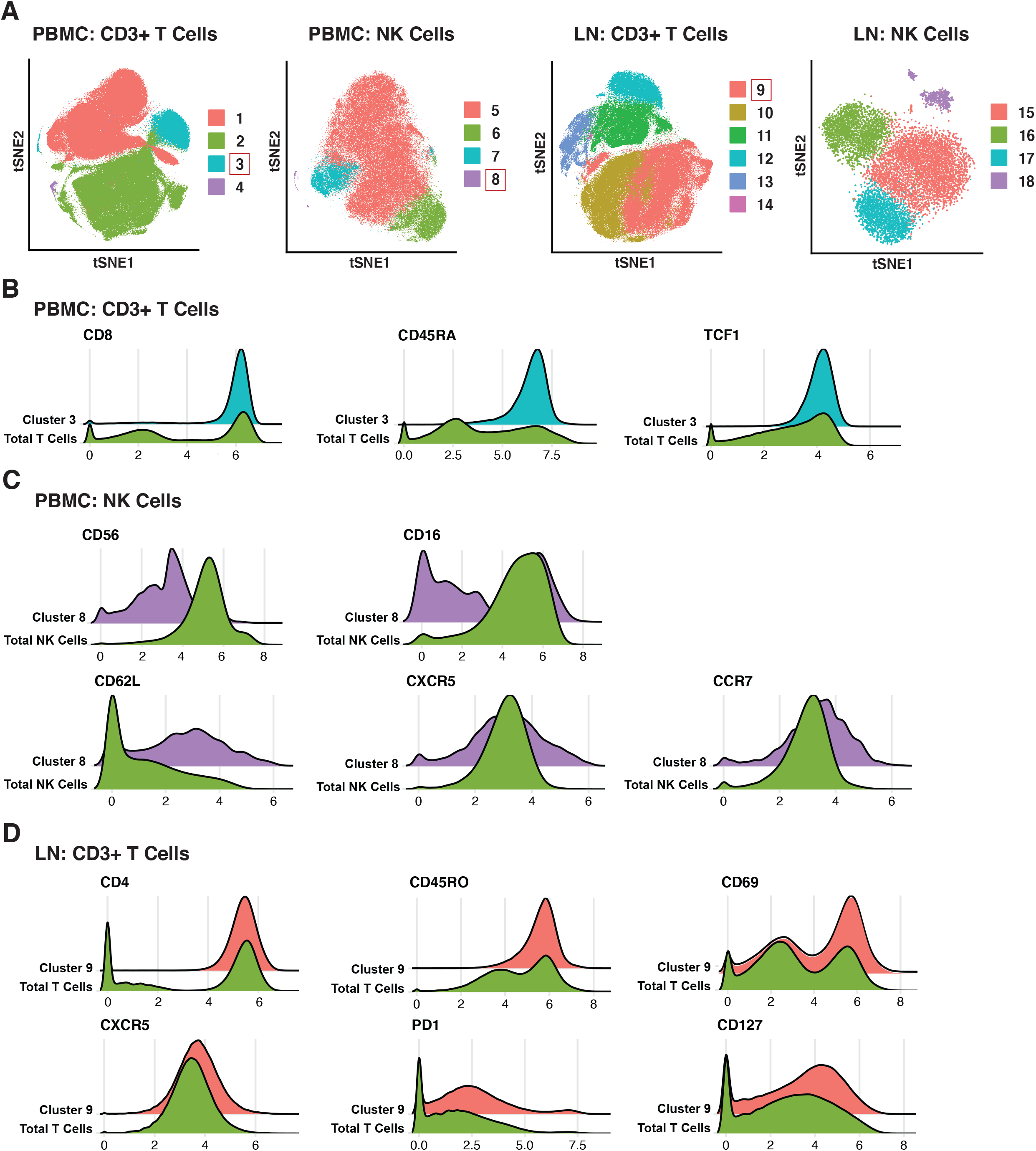
Clusters of T and NK cells associated with acutely treatment of HIV. **(A)** tSNE visualization of T and NK cell clusters from PBMCs and LNs. Clusters significantly enriched in the acutely-treated group are boxed in red. **(B)** Cluster 3 of blood T cells was enriched in the acutely-treated group relative to controllers (P < 0.0001) and the chronically-treated group (P < 0.009). Cells in Cluster 3 expressed relatively high levels of CD8, CD45RA, CD27, CCR7, and TCF1. **(C)** Cluster 8 of blood NK cells was enriched in the acutely-treated group relative to chronically-treated group (P < 0.05). Cells in Cluster 8 expressed low levels of CD56, and approximately half of them were CD16+. Cluster 8 also included cells expressing high levels of lymph node-homing receptors CD62L, CCR7, and CXCR5. **(D)** Cluster 9 of LN T cells was enriched in the acutely-treated group relative to controllers (P < 0.03) and the chronically-treated group (P < 0.0001). Cluster 9 corresponded to CD4+ T cells expressing high levels of the T resident memory (Trm) marker CD69, the T follicular helper (Tfh) markers PD1 and CXCR5, and the alpha chain of the IL7 receptor CD127. A generalized linear mixed model (GLMM) was used to estimate the association between cluster membership and group (controllers, acutely-treated, or chronically-treated). The log odds ratio was retrieved from the coefficient estimate of each variable. All p-values were adjusted using the FDR, and a threshold of 0.05 was used to determine significance.

We then searched for unique features and expression profiles in the clusters that were over-represented among the acutely-treated group (Fig. 2B-D; expression levels of all antigens from all clusters are shown in Fig. S2-5). Cluster 3 of blood T cells expressed relatively high levels of CD8, CD45RA, and TCF1 (Fig. 2B), consistent with a more Tn-like phenotype (Fig. 1C). Cluster 8 of blood NK cells preferentially expressed low levels of CD56 and included both CD16- and CD16+ cells (Fig. 2C), the latter of which are typically rare but expanded in PLWH, and are associated with HIV control (Bradley et al., 2018). Cluster 8 also expressed relatively high levels of the lymph node homing receptors CD62L and CCR7, and harbored the cells expressing the highest levels of CXCR5, suggesting these cells may be poised to migrate into the follicles of LNs. Within the LNs, Cluster 9 of T cells expressed relatively high levels of CD4, CD45RO, CD69, suggesting a T resident memory phenotype; CXCR5 and PD1, suggesting a Tfh-like phenotype; and CD127 (the alpha chain of the IL7 receptor), suggesting homeostatic proliferation potential of these cells. In sum, these data suggest acutely-treated individuals are enriched for subsets of T and NK cell exhibiting distinguishing phenotypic features.

### 2.4 Blood-derived T Cells From HIV+ Individuals Are More Activated Than Those from Lymph Nodes

Having defined clusters of lymphocytes that associate with a specific state of HIV control, we next conducted comparisons between the paired PBMC and LN specimens to identify differences due to cell origin. As the overall phenotypes of T cells were overall similar between individuals (Fig. 1A, 1B, 3A *right*) we combined the data from the three patient groups together for this analysis. This analysis suggested lymphocytes from LNs to be profoundly different from their blood counterparts in that the two types of cells occupied very distinct regions of tSNE (Fig. 3A *left*). The major subsets (Tn; Tm; Tcm; Tem; Tfh) of CD4+ (Fig. 3B) and CD8+ (Fig. 3C) T cells also demonstrated overall differences between blood (in grey) and LNs (in color). Manual gating also revealed different frequencies of these subsets between blood and LNs, in particular among the CD4+ Tem, CD8+ Tn, CD8+ Tm, and CD8+ Tem subsets (Fig. 3D-E).

**Figure 3.**
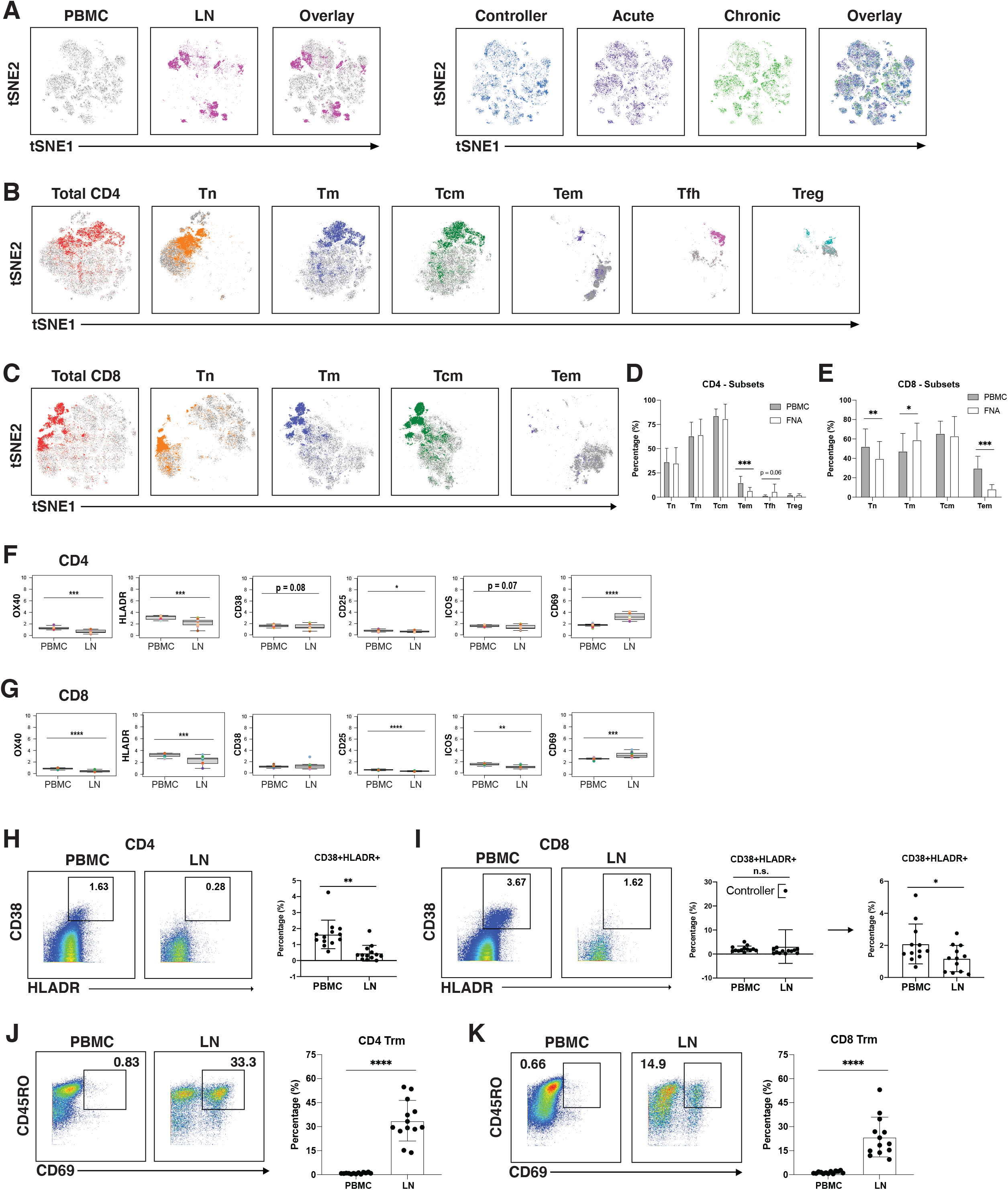
T cells from blood of PLWH are more activated than those from lymph nodes. **(A)** tSNE plots demonstrating profound differences between T cells from blood (grey) and LNs (colored; left), and more subtle differences between patient groups (controllers in blue, acutely-treated in purple, and chronically-treated in green; right). **(B)** tSNE plots depicting phenotypic differences among subsets of CD4+ T cells from blood (grey) vs. LNs (colored). **(C)** tSNE plots depicting phenotypic differences among subsets of CD8+ T cells from blood (grey) vs. LNs (colored). CD4+ Tem **(D)** and CD8+ Tem and CD8+ Tn **(E)** are more frequent in blood than in LNs. Shown are frequencies of T cell subsets relative to total PBMC (grey bars) or LNs (white bars), displayed as bar graphs. Cell subsets are as described in Fig. 1C. CD4+ **(F)** and CD8+ **(G)** T cells from blood express high levels of activation markers, with the exception of activation/tissue residency marker CD69. Shown are the mean signal intensity (MSI) levels of the indicated activation markers assessed in T cells from PBMCs and LNs. Data are displayed as bar graphs with individual donors’ values as colored dots. The remaining antigens are presented in **Fig. S6-S7**. Activated CD38+HLADR+ CD4+ **(H)** and CD8+ **(I)** T cells are more frequent in PBMCs than LNs. Gates for activation of T cells dually expressing the two activation markers CD38 and HLA-DR (left). The bar graphs (right) show the percentages of CD38+HLA-DR+ T cells in PBMCs and LNs among all donors examined. For CD8+ T cells, one extreme outlier proved to be an HIV controller; data analyzed after removing this outlier are shown on the far right. **(J)** CD4+ and **(K)** CD8+ T resident memory (Trm) cells are enriched in LNs. Trm cells were defined as those dually expressing CD45RO and CD69 (left). Their frequencies in PBMCs and LNs are depicted as bar graphs (right). *p<0.05, **p<0.01, ***p<0.001, ****p<0.0001 as determined using the Student’s paired t test and adjusted for multiple testing using the Benjamini-Hochberg for FDR.

Interestingly, activation states differed between T cells from blood vs. LNs. When we examined the mean signal intensity (MSI) of the antigens quantitated by CyTOF (Fig. S6-7), we found the activation markers OX40, HLADR, and CD25 to be expressed at significantly higher levels on both CD4+ and CD8+ T cells from PBMCs (Fig. 3F-G). Blood T cells also expressed higher levels of the activation marker ICOS, but this did not reach statistical significance for the CD4 subset (Fig. 3F-G). Expression of the activation marker CD38 trended higher on blood than LN CD4+ T cells, albeit not significantly (Fig. 3F). To confirm the elevated activation status of the blood T cells, we manually gated on T cells dually expressing CD38 and HLADR (Fig. 3H-I), as these two markers have been used to define T cell activation state during HIV infection (Kestens et al., 1992; Doisne et al., 2004; Papagno et al., 2004). CD38+HLADR+ CD4+ T cells were significantly more frequent in PBMCs than LNs (Fig. 3H), while for CD8+ T cells, no significant differences were observed (Fig. 3I). However, removal of one outlier individual, a controller with unusually high proportions of these cells in LNs, lead to significant differences in the CD8+ T cell compartment as well (Fig. 3I).

In contrast to the aforementioned activation markers, CD69 was increased on T cells from LNs as compared to T cells from PBMCs (Fig. 3F-G). This is likely because while an activation marker in blood cells, CD69 is also the canonical marker for tissue resident memory T cells (Trm) (Baeyens et al., 2015). Manual gating of Trm (CD45RO+CD69+) cells confirmed that the frequency of CD4+ (Fig. 3J) and CD8+ (Fig. 3K) Trm was significantly greater in LNs than in PBMCs. Therefore, with the exception of CD69, activation markers were elevated on blood as compared to LN T cells.

### 2.5 Memory CD8+ T Cells Expressing CXCR5 Are Enriched in Lymph Nodes and Exhibit a Long-Lived, Inhibited Phenotype

As CXCR5 expression can direct CD8+ T cells to the follicles, a major site of HIV persistence (Ayala et al., 2017; Haran et al., 2018), we next determined to what extent CXCR5 was expressed on CD8+ Tm cells from blood and LNs. In addition to examining total Tm cells, we also examined Trm cells, the most abundant T cell subset in human tissues (Thome et al., 2014). The percentage of Tm and Trm cells expressing CXCR5 was higher in LNs than in blood (Fig. 4A-B). To gain a better understanding of the features of these follicular CXCR5-expressing CD8+ T cells, we compared their expression levels of homing receptors and activation/differentiation markers to that of their CXCR5-counterparts. Expression of tissue residency marker CD69 was increased in CXCR5+ cells (Fig. 4C), albeit insignificantly. Expression of tissue homing receptors CCR6 and CXCR4 was significantly increased on CXCR5+ cells, as was expression of the lymph node homing receptors CCR7 and CD62L (Fig. 4C), suggesting the ability of these cells to migrate into multiple lymphoid tissues. TCF1 and CD127, associated with long-term survival of T cells, were also expressed at elevated levels on the CXCR5+ cells. As NK cell receptors have previously been reported to be expressed on T cells and to regulate their function (Cibrian and Sanchez-Madrid, 2017), we also assessed their expression on CXCR5+ vs. CXCR5-CD8+ Tm cells (Fig. 4C). Surprisingly, the inhibitory NK cell receptors KIR2DL1, CD94, and NKG2A were significantly increased on the CXCR5+ cells, while the activating NK cell receptor NKG2D was decreased albeit insignificantly. These results suggest that within lymph nodes, CXCR5+ CD8+ T cells, relative to their CXCR5-counterparts, have broad tissue homing capabilities and are long-lived, but may be restrained in their effector functions as they express multiple markers associated with suppression of effector functions. Whether CXCR5+ NK cells exhibit similar phenotypic features could not be reliably assessed as there were too few of these cells in our specimens.

**Figure 4.**
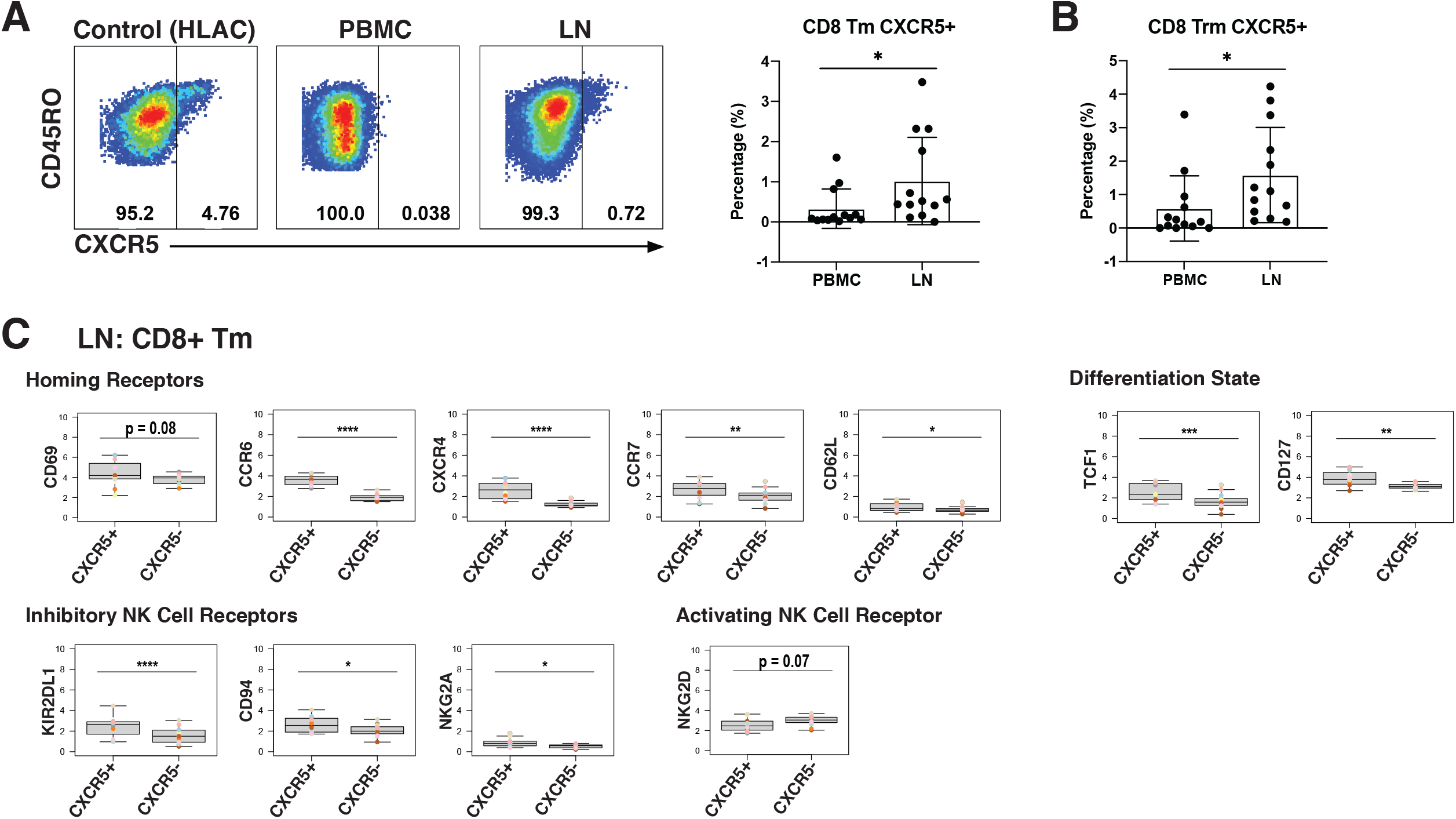
LNs from PLWH are enriched for CXCR5+ CD8+ T cells expressing high levels of inhibitory NK cell receptors. **(A)** CXCR5+ memory T cells are more frequent in LNs than in blood. Gating strategy (left) and frequencies of CXCR5+ cells (right) among memory CD8+ T cells from paired PBMCs and LNs. The gating strategy for CXCR5+ cells was established using human lymphoid aggregate cultures (HLAC) of tonsils, which harbor a distinct population of these cells. **(B)** CXCR5+ tissue-resident memory T cells (Trm) are more frequent in LNs than in blood. **(C)** Lymphoid CXCR5+ memory CD8+ T cells express higher levels of tissue homing receptors, markers of long-lived memory cells, and inhibitory NK receptors, relative to their CXCR5- counterparts do. Shown are MSIs of the indicated homing receptors, differentiation state markers, inhibitory and activating NK cell receptors. *p<0.05, **p<0.01, ***p<0.001, ****p<0.0001 as assessed using the Student’s paired t test and adjusted for multiple testing using the Benjamini-Hochberg for FDR.

### 2.6 Memory-like NK Cells Are Enriched in LNs and Express a Distinct Repertoire of Activating NK Receptors Relative to Non-Memory NK Cells

Finally, we assessed for phenotypic differences between blood- and LN-derived NK cells. NK cells from blood vs. LN were concentrated within different areas of the tSNE (Fig. 5A), and this was true of the 3 different subsets (CD56^bright^, CD56^dim^, CD16-) of NK cells (Fig. 5B). Furthermore, LNs harbored more CD56^bright^ and fewer CD56^dim^ NK cells than PBMCs did, but the same percentage of CD56-NK cells (Fig. 5C).

**Figure 5.**
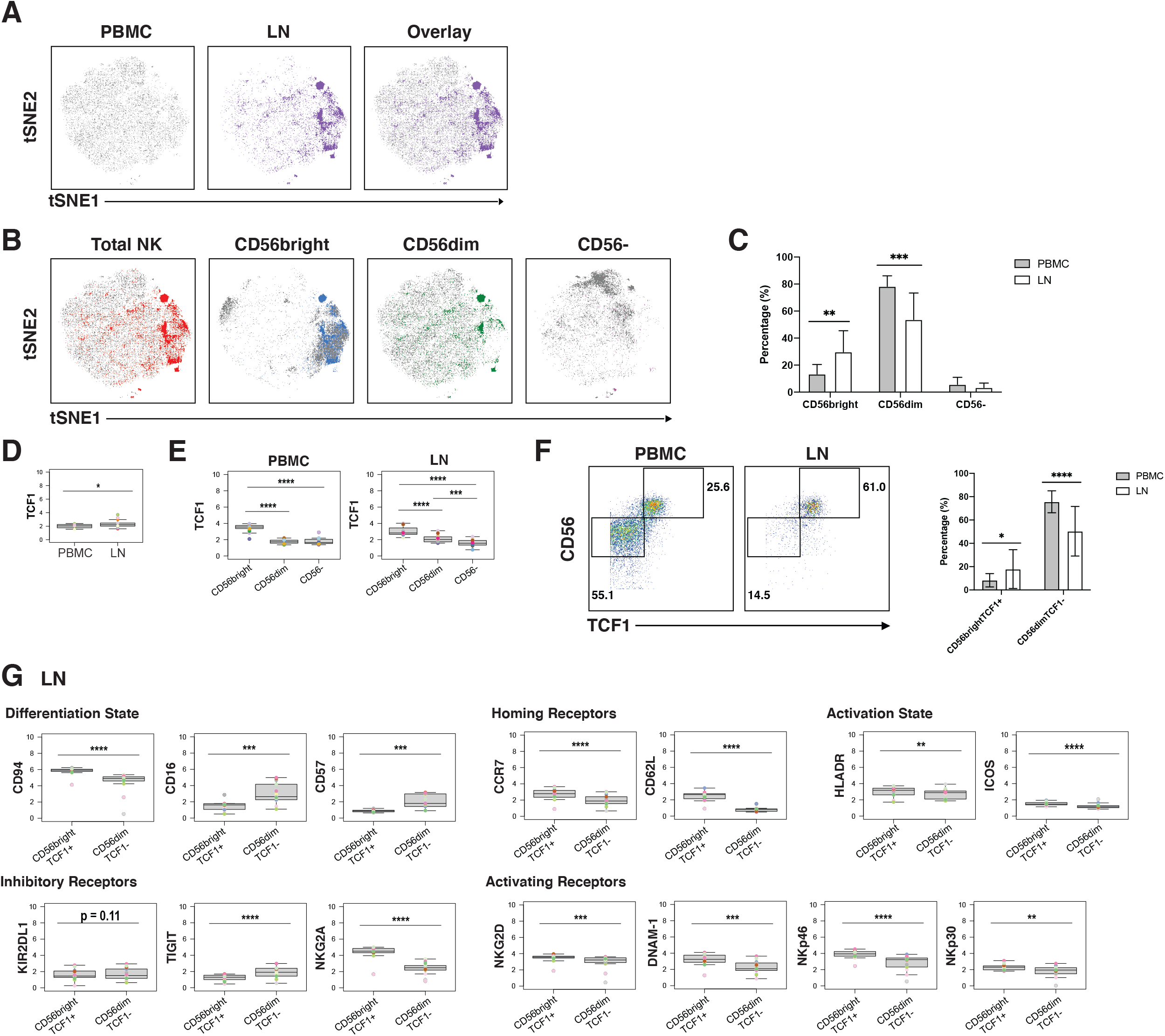
CD56^bright^ NK cells from PLWH display phenotypic features of memory and are enriched in LNs. **(A)** tSNE plots demonstrating the overall differences between total NK cells from blood (grey) and LN (colored). **(B)** NK cell subsets, defined by CD56 expression (total, CD56^bright^, CD56^dim^, CD56-), are phenotypically distinct in blood (grey) as compared to LNs (colored). **(C)** NK cells from PBMCs have increased frequencies of CD56^dim^ cells, while NK cells from LNs have increased frequencies of CD56^bright^ cells. Shown are the percentages of the indicated populations among total NK cells. **(D)** TCF1 expression is higher in NK cells from LNs than from blood. MSI of TCF1 in total NK cells from PBMCs and LNs are shown. Expression levels of other antigens on these two cell populations are presented in **Fig. S8. (E)** TCF1 expression is highest in the CD56^bright^ subset of NK cells. MSI of TCF1 in NK cell subsets from PBMCs and LNs are shown. **(F)** LNs harbor a larger proportion of CD56^bright^TCF1+ NK cells than do PBMCs. Representative gates (left) and frequencies (right) of CD56^bright^TCF1+ and CD56^dim^TCF1-NK cells in PBMCs and LNs. **(G)** LN CD56^bright^TCF1+ NK cells are less differentiated and express increased levels of LN homing receptors, activation markers, and activating NK receptors than LN CD56^dim^TCF1-NK cells. Shown are MSI levels of the indicated markers assessed in CD56^bright^TCF1+ and CD56^dim^TCF1- NK cells from LNs. Plots comparing MSI of the indicated markers between CD56^bright^TCF1+ and CD56^dim^TCF1-NK cells from PBMCs are presented in **Fig. S8**. *p<0.05, **p<0.01, ***p<0.001, ****p<0.0001 as assessed using the Student’s paired t test and for MSIs, adjusted for multiple testing using the Benjamini-Hochberg for FDR.

To further assess the differences between NK cells derived from blood vs. LN, we compared the MSI for all antigens from our CyTOF panel (Fig. S8A). This analysis revealed that TCF1 was significantly increased in LN relative to blood NK cells (Fig. 5D). Interestingly, *TCF7* (transcript for TCF1) marks a memory-like NK cell subset within the blood of PLWH, and this subset was hypothesized to contribute to HIV control (Wang et al., 2020). As this memory-like NK cell subset was also shown to be CD56^bright^ (Wang et al., 2020), we then compared MSI expression of TCF1 between the 3 major NK cell subsets (CD56^bright^, CD56^dim^, CD16-). We found, as expected (Wang et al., 2020), TCF1 expression levels to be highest in the CD56^bright^ subset in blood, and further demonstrated that this was true in the LN compartment as well (Fig. 5E). Manual gating revealed that the LN harbored significantly higher proportions of these memory-like CD56^bright^TCF1+ NK cells, while blood harbored more CD56^dim^TCF1-NK cells (Fig. 5F).

To better define the CD56^bright^TCF1+ NK cells in LNs, we compared the MSI of markers of differentiation and activation states, homing receptors, and inhibitory and activating NK cell receptors between the CD56^bright^TCF1+ (memory-like) vs. CD56^dim^TCF1-(not memory-like) NK cells (Fig. 5G). LN CD56^bright^TCF1+ NK cells expressed high levels of the co-receptor CD94, an antigen recently reported to be expressed on CD56^bright^TCF1+ NK cells from the blood of PLWH (Wang et al., 2020). They also expressed high levels of the LN homing receptors CCR7 and CD62L, high levels of the activation markers HLADR and ICOS, and low levels of the NK cell differentiation markers CD16 and CD57, suggesting that they may be less terminally differentiated (Dogra et al., 2020). They also expressed high levels of the inhibitory receptor NKG2A, and high levels of the activation receptors NKG2D, DNAM-1, NKp46, and NKp30, relative to LN CD56^dim^TCF1-NK cells. By contrast, low levels of the inhibitory receptor TIGIT were observed in these cells, and the inhibitory NK cell receptor KIR2DL1 was also decreased, albeit not significantly. Interestingly, the differences observed between CD56^bright^TCF1+ and CD56^dim^TCF1-NK cells from the LNs were recapitulated within the blood compartment (Fig. S8B).

In sum, these results show that a memory-like CD56^bright^ TCF1+ NK cell subset previously described in the blood of PLWH is enriched in LNs, and that these cells express high levels of multiple activation markers and activating NK cell receptors, suggesting they exist in a state of heightened activation relative to non-memory (CD56^dim^TCF1-) NK cells.

## 3 Discussion

In this study, we used high-parameter single-cell phenotyping by CyTOF to define the features of blood- and LN-derived T and NK cells from three groups of aviremic PLWH: elite controllers, and virally-suppressed non-controllers who had initiated ART during the acute vs. chronic phases of infection. Our main findings are that T and NK cells from the acutely-treated individuals harbour unique phenotypic features, and differ markedly depending on whether they derived from the blood or LN compartment.

Three clusters of cells were significantly enriched in the acutely-treated individuals. One of these (Cluster 3) corresponded to Tn cells. The second, Cluster 8, corresponded to NK cells that express high levels of the homing receptors CD62L, CCR7, and CXCR5, suggesting the ability of these cells to migrate into or be retained within the LN follicle. Interestingly, Cluster 8 included the CD56-CD16+ subset of NK cells enriched within HIV+ individuals who produce broadly neutralizing antibodies (bnAbs) against HIV (Bradley et al., 2018), suggesting a role for these NK cells in working together with bnAbs to control HIV replication. How these NK cells may do this is unclear, but it may reflect the fact that, unlike CD56+ NK cells, they are unable to restrict the stimulation of antibody production by CD4+ T cells (Bradley et al., 2018). Consistent with this hypothesis, Cluster 9, which like Cluster 8 was over-represented in the acutely-treated group, consisted of LN CD4+ T cells that exhibited features of Tfh, which are known to be important for providing help to B cells, and that expressed high levels of CD127, which promotes T cell survival and proliferation (Kaech et al., 2003; Kondrack et al., 2003). Because individuals who initiated ART during the acute phase of infection are more likely to become post-treatment controllers (PTCs) upon treatment interruption (ATI) (Lodi et al., 2012; Saez-Cirion et al., 2013; Maenza et al., 2015; Sneller et al., 2017), future studies should determine if CD56-CD16+ NK cells and CD127-expressing Tfh-like CD4+ T cells are enriched in PTCs, as this would suggest a causal role for these cells in viremic control after ART withdrawal.

In-depth comparisons of CD8+ T cells from the blood and tissue compartments from our cohort revealed that those from LNs were relatively quiescent, and preferentially expressed CXCR5. As CXCR5+ CD8+ T cells are poised to enter the LN follicle where much of the HIV reservoir resides (Brenchley et al., 2012; Lindqvist et al., 2012; Perreau et al., 2013; Banga et al., 2016), they have the potential to control viral replication. However, we found that these cells expressed elevated levels of a number of inhibitory NK cell receptors, including KIR2DL1, NKG2A, and NKG2A’s co-receptor CD94, which can dampen CD8+ T cells’ effector function (Moser et al., 2002; Nattermann et al., 2006; Sullivan et al., 2021). Our finding that expression of these receptors is elevated on LN CD8+ T cells, suggests that within the lymphoid compartment these cells may be preferentially restricted in their effector functions. Consistent with this notion are recent reports that LN HIV-specific CXCR5+ CD8+ T cells from both controller and non-controller PLWH exhibit poor cytolytic activity *ex vivo* (Reuter et al., 2017; Nguyen et al., 2019). Therefore, strategies aimed to eradicate the lymphoid tissue HIV reservoir may need to not only increase CD8+ T cell entry into the B cell follicle, but also release the intrinsic breaks on their cytolytic or other effector mechanisms, perhaps through antagonism of these inhibitory receptors.

There was also an enrichment of a unique subset of memory-like NK cells in LNs of PLWH. These cells, identified as CD56^bright^TCF1+ NK cells, are phenotypically similar to memory-like NK cells previously described in the blood of PLWH (Wang et al., 2020). In both the blood and LN compartments, the CD56^bright^TCF1+ NK cells exhibited decreased expression of NK cell maturation markers CD16 and CD57, and increased expression of the homing receptors CCR7 and CD62L, relative to their CD56^dim^TCF1-counterparts. They also expressed increased levels of the activation markers HLADR and ICOS, increased expression of the NKG2A/C co-receptor CD94, and activating receptors NKG2D, DNAM1, NKp46, and NKp30. These results suggest overall phenotypic similarities between these memory-like NK cells in the blood and LN compartments. Interestingly, memory-like CD56^bright^ NK cells were recently described in BLT humanized mice vaccinated with HIV envelope protein, and these cells, like the ones we have characterized, also expressed decreased levels of CD16 and CD57, and increased levels of CD62L and NKG2A (Nikzad et al., 2019). As these BLT tissue-derived memory-like NK cells were capable of mediating antigen-specific recall responses *ex vivo* (Nikzad et al., 2019), the LN CD56^bright^TCF1+ NK we characterized may also be capable of mediating such recall responses and play an active role in controlling HIV replication within tissues. In summary, memory-like CD56^bright^TCF1+ NK cells are enriched in the lymphoid compartment and exhibit similar phenotypic features as their blood counterparts, including expression of numerous NK activating receptors which may aid in their ability to respond to and kill HIV-infected cells.

Our study has limitations. Our sample size was small and the findings reported here should be validated in larger cohorts. Additionally, due to the limited number of cells available (particularly from LNs), we were unable to evaluate functional responses of T and NK cells. Future studies should define how the blood and lymphoid T and NK cell subsets characterized here respond to HIV-infected target cells. Despite these limitations, our data strongly suggest that the phenotypic features of T and NK cells are affected by the timing of when ART is initiated during HIV infection, and whether these cells are isolated from the blood or LNs. We find that CXCR5-expressing CD8+ T cells and memory-like NK cells are both enriched within the LNs of PLWH. These effector cells may be good candidates to unleash for immune-based therapeutic approaches to achieve ART-free HIV remission.

## 5 Author Contributions

A.F.G. designed and conducted experiments, conducted data analysis, and wrote the manuscript. X.L. wrote scripts for data analysis. J.N. provided technical advice. S.G.D. established the SCOPE cohort. R.H. conducted SCOPE participant enrollment, S.G.D., and S.A.L. established the UCSF Acute HIV study within the SCOPE cohort, S.A.L. conducted the acute HIV participant enrollment, and P.V. conducted specimen collection. M-G.S. and R.T. provided bioinformatics assistance. C.A.B. provided technical advice. W.C.G. supervised data analysis and edited the manuscript. S.A.L. and N.R.R. conceived ideas and S.G.D., W.G., S.A.L, and N.R.R. obtained funding to support the study. N.R.R. designed experiments, edited the manuscript, and supervised the study. All authors read and approved the manuscript.

## 7 Acknowledgments

We acknowledge NIH DRC Center grant P30 DK063720 and S10 1S10OD018040-01 for use of the CyTOF instrument. We also thank S Tamaki, TK Peech, and C Bispo for CyTOF assistance at the Parnassus Flow Core; V Tai and M Kerbleski for assistance with the SCOPE specimens; F Chanut for editorial assistance; and R Givens for administrative assistance.

## 9 Supplementary Figure Legends

**Figure S1.**
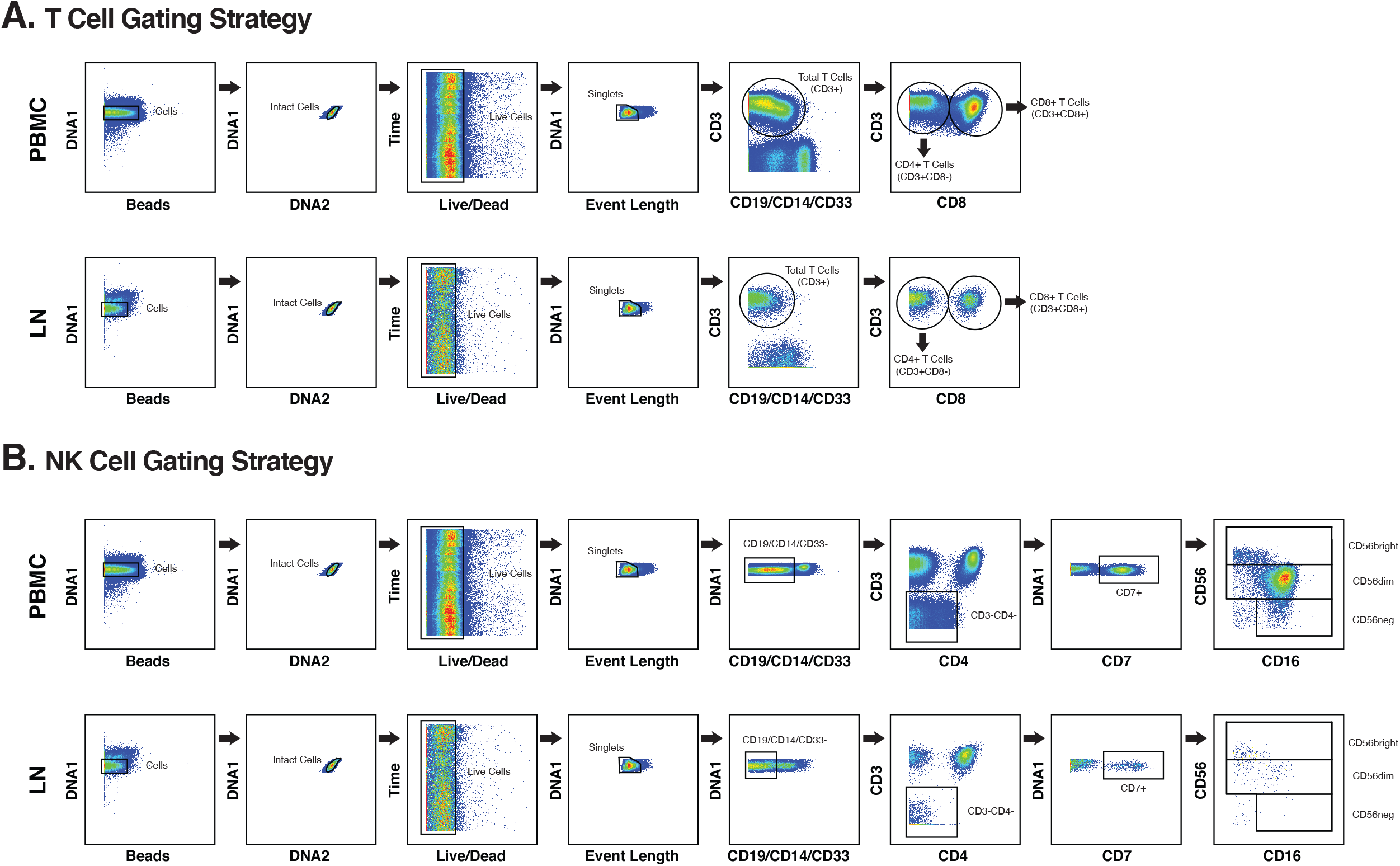
CyTOF gating strategy to identify total T cells, CD4+ T cells, CD8+ T cells, and NK cells from PBMCs and LNs of PLWH. **(A)** Shown are gating strategies to identify live, singlet, total T cells, CD4+ T cells, and CD8+ T cells from a representative PBMC and LN specimen. (B) Shown are gating strategies to identify live, singlet NK cells from a representative PBMC and LN specimen.

**Figure S2.**
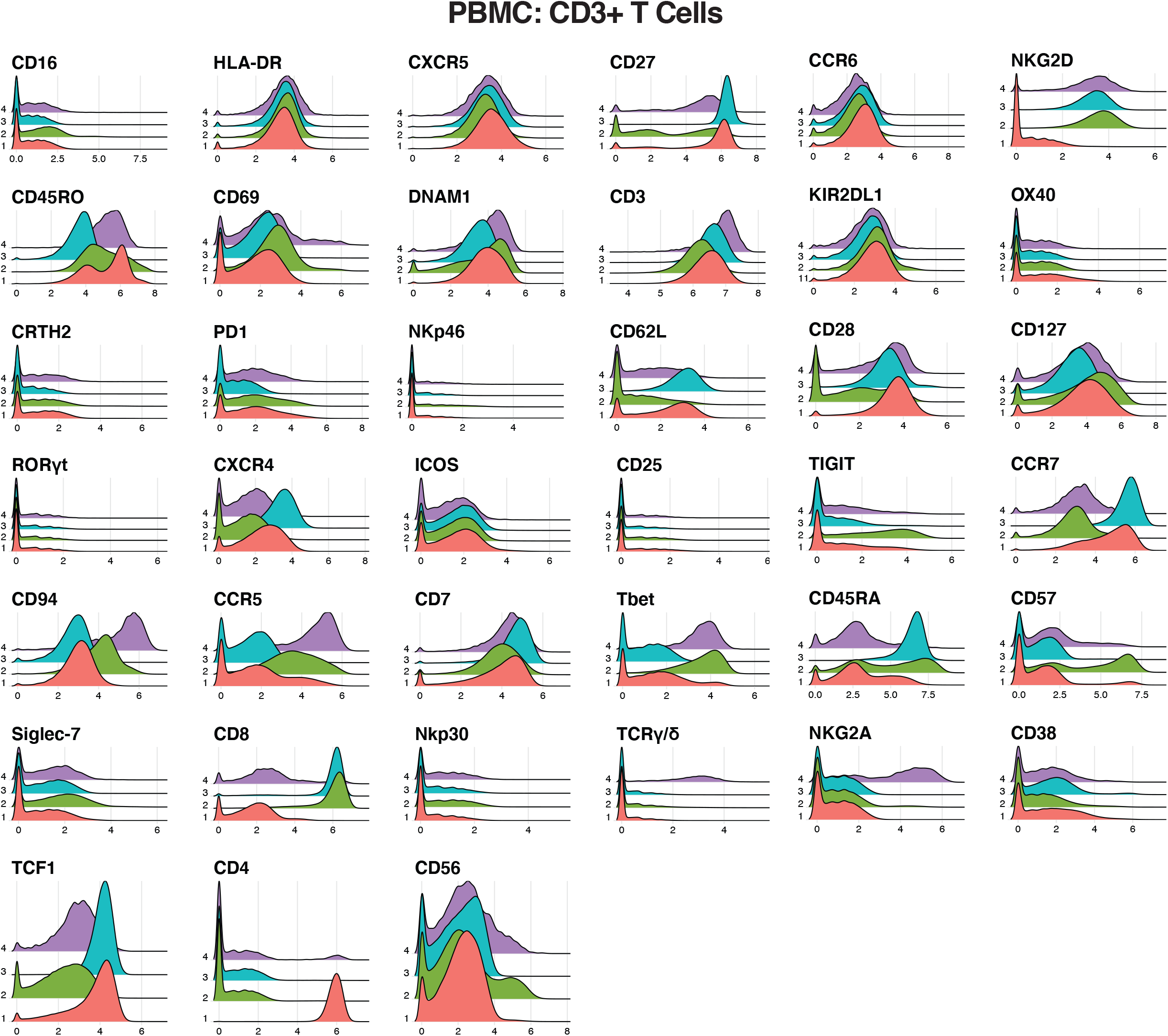
Expression levels of antigens on Seurat-identified clusters of T cells from PBMCs of PLWH. Shown are histogram plots depicting expression levels of the indicated antigens on PBMC T cells (live, singlet, CD19-CD14-CD33-CD3+) cells from clusters identified in Fig. 2A. For a detailed explanation on how the datasets were clustered, see Materials and Methods.

**Figure S3.**
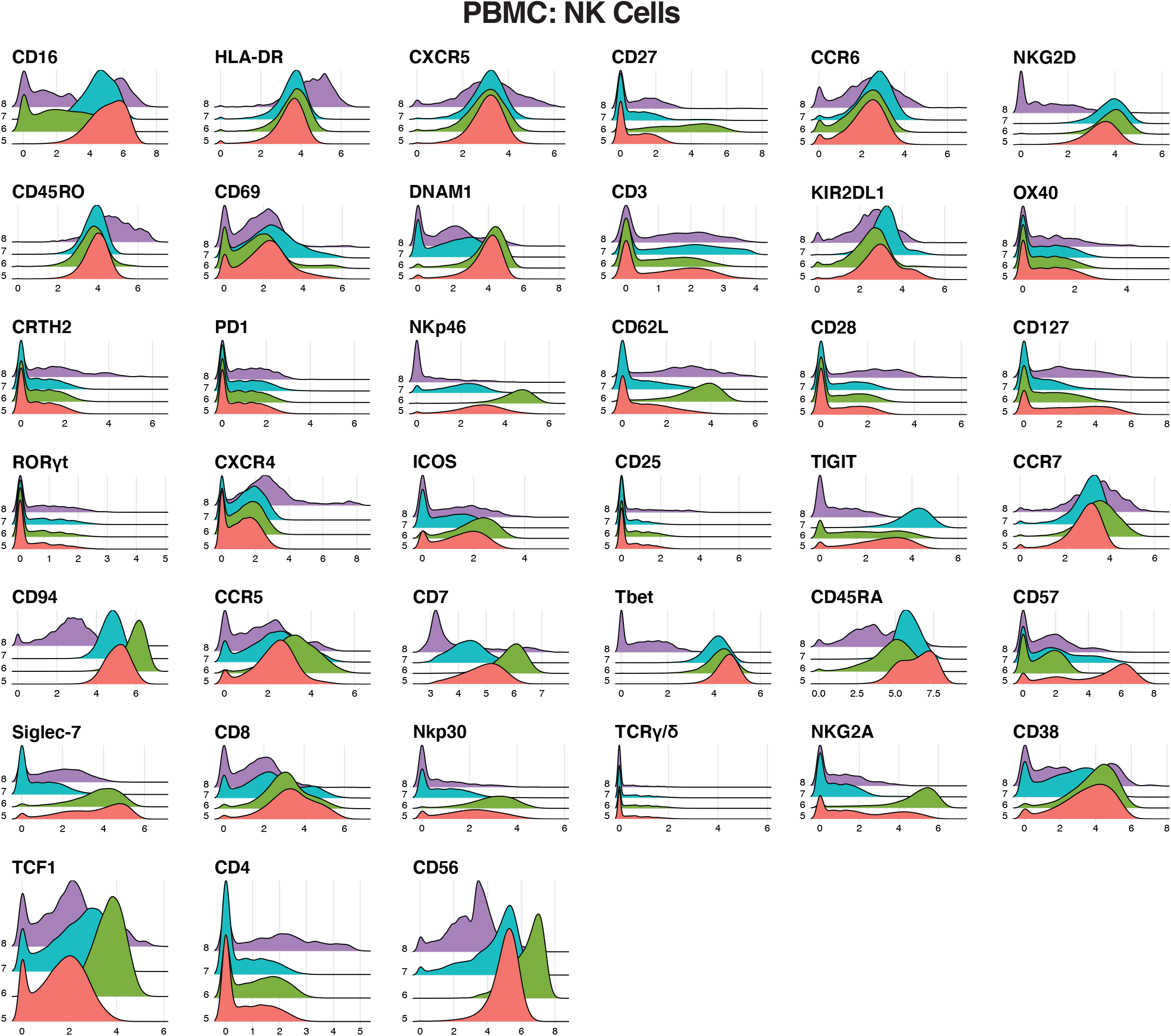
Expression levels of antigens on Seurat-identified clusters of NK cells from PBMCs of PLWH. Expression levels of the indicated antigens on PBMC NK cells (CD19-CD14-CD33-CD3-CD4-CD7+) from clusters identified in Fig. 2A are depicted as histogram plots.

**Figure S4.**
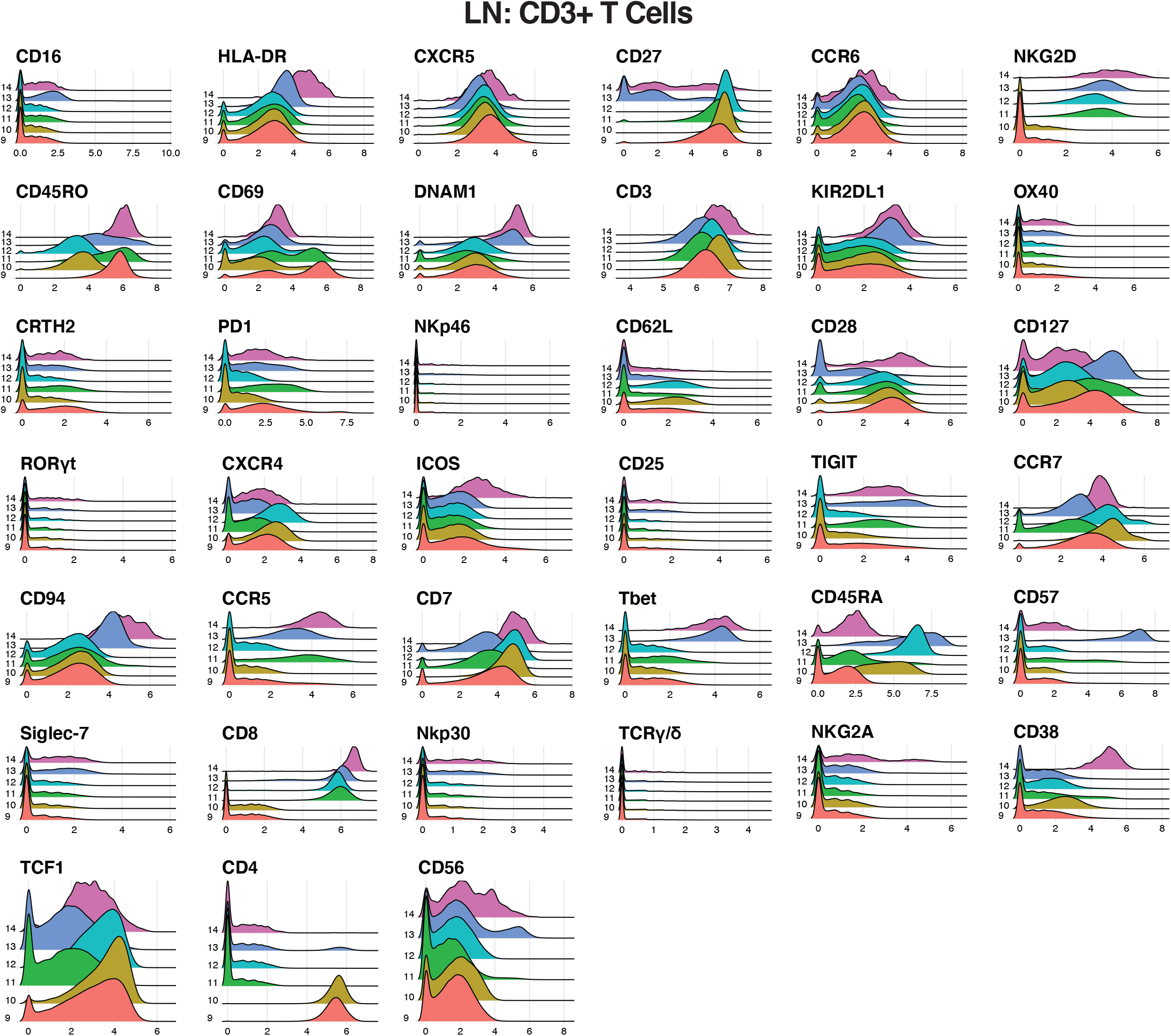
Expression levels of antigens on Seurat-identified clusters of T cells from LNs. Expression levels of the indicated antigens on LN T cells (live, singlet, CD19-CD14-CD33-CD3+) from clusters identified in Fig. 2A are depicted as histogram plots.

**Figure S5.**
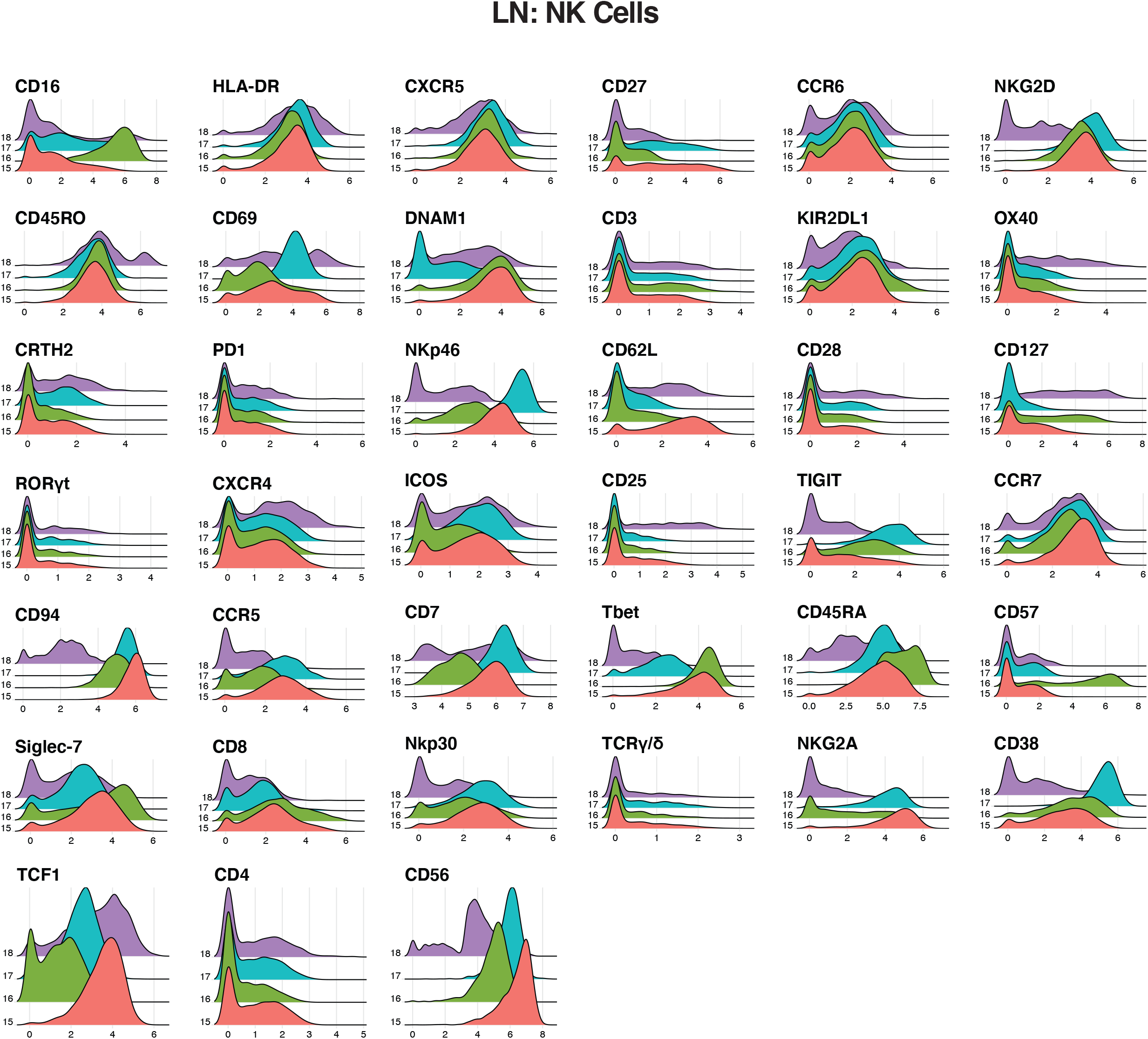
Expression levels of antigens on Seurat-identified clusters of NK cells from LNs. Expression levels of the indicated antigens on LN NK cells (CD19-CD14-CD33-CD3-CD4-CD7+) from clusters identified in Fig. 2A are depicted as histogram plots.

**Figure S6.**
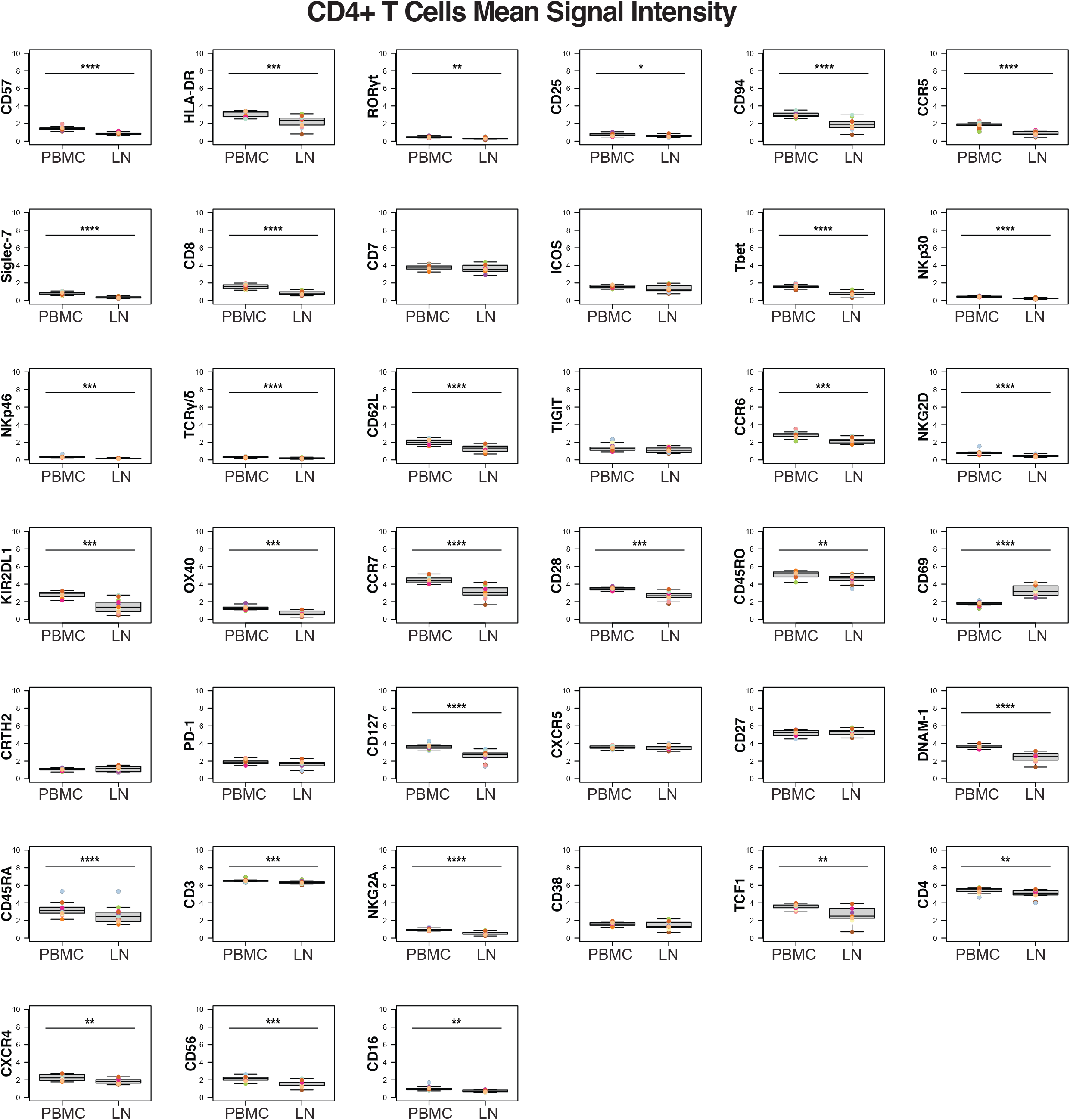
Expression levels of 39 surface and intracellular antigens in CD4+ T cells from PBMCs and LNs of PLWH. Shown are the mean signal intensity (MSI) values of CD4+ T cells from PBMCs as compared to LNs. *p<0.05, **p<0.01, ***p<0.001, ****p<0.0001 as assessed using the Student’s unpaired t-test and adjusted for multiple testing using the Benjamini-Hochberg for FDR.

**Figure S7.**
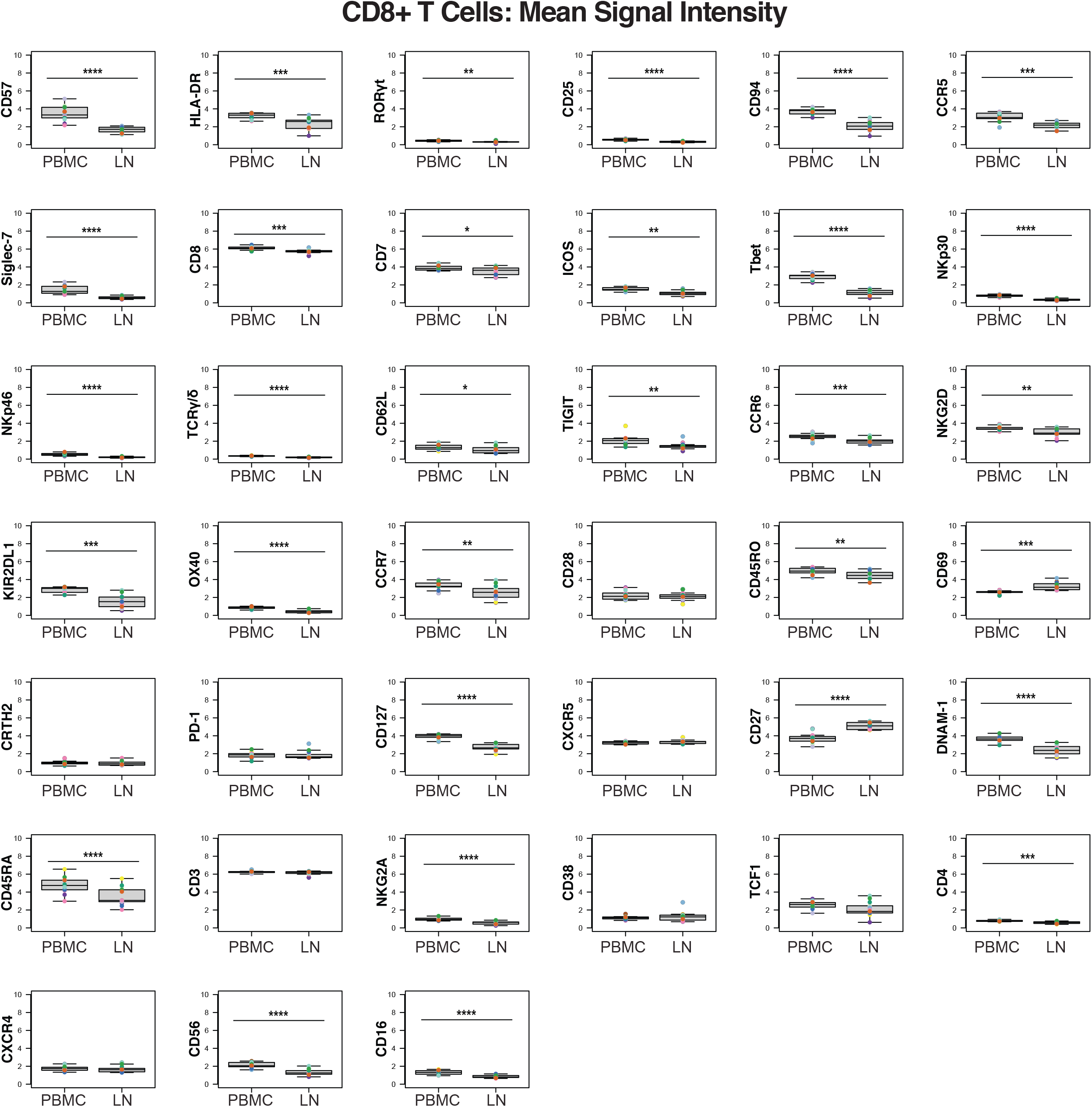
Expression levels of 39 surface and intracellular antigens in CD8+ T cells from PBMCs and LNs of PLWH. Shown are the mean signal intensity (MSI) values of CD8+ T cells from PBMCs as compared to LNs. *p<0.05, **p<0.01, ***p<0.001, ****p<0.0001 as assessed using the Student’s unpaired t-test and adjusted for multiple testing using the Benjamini-Hochberg for FDR.

**Figure S8.**
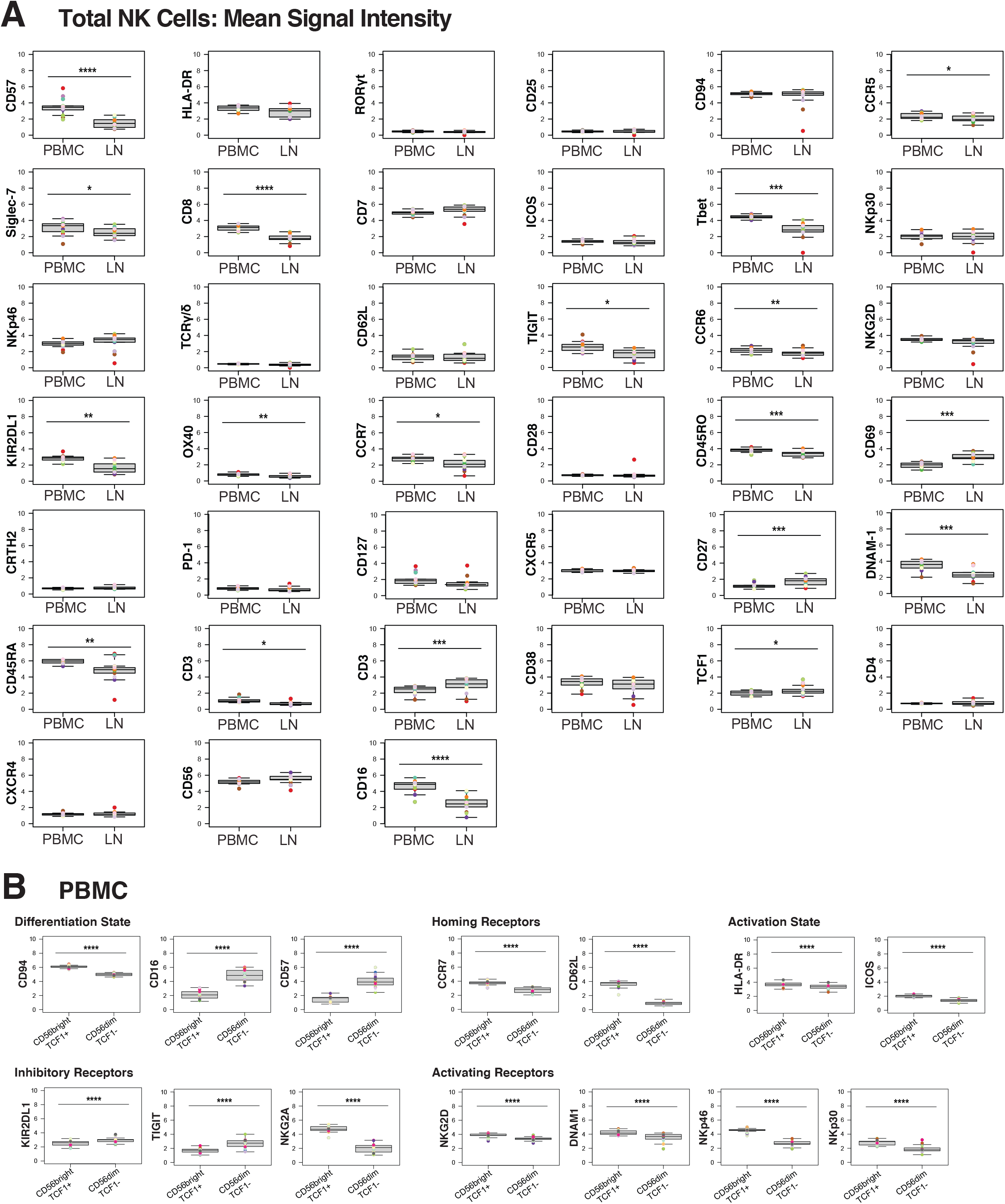
Expression levels of 39 surface and intracellular antigens in NK cells from PBMCs and LNs of PLWH. **(A, B)** Mean signal intensity (MSI) values comparing NK cells from PBMCs vs. LNs (A), or between CD56^bright^TCF1+ and CD56^dim^TCF1-NK cells from PBMCs (B). PBMC CD56^bright^TCF1+ cells are less differentiated and express higher levels of LN homing receptors, activation markers, and activating NK receptors than do PBMC CD56^dim^TCF1-NK cells. *p<0.05, **p<0.01, ***p<0.001, ****p<0.0001 as determined using the Student’s unpaired t-test and adjusted for multiple testing using the Benjamini-Hochberg for FDR.

## 1 Supplementary Tables

**Table S1.** Participant characteristics. The age, gender, race/ethnicity, antiretroviral regimen (ART), viral load (VL) levels in copies/ml, absolute CD4 T cell count, and absolute CD8 T cell count of each participant at time of specimen collection are indicated.

| PID | Category | Age | Gender | Race/Ethnicity | ART | VL | CD4 Count | CD8 Count |
| --- | --- | --- | --- | --- | --- | --- | --- | --- |
| 1349* | Controller | 63 | Male | White | None | <40 | 573 | 1042 |
| 1508* | Controller | 56 | Male | Black/African American | None | 81 | 425 | 1394 |
| 1559* | Controller | 60 | Male | Black/African American | None | <40 | 491 | 442 |
| 1775* | Controller | 39 | Male | White | None | 776 | 851 | 882 |
| 3736 | Controller | 60 | Male | White | None | <40 | 853 | 406 |
| 8007* | Acutely-treated | 66 | Male | White | BIC/FTC/TAF | <40 | 702 | 646 |
| 8009* | Acutely-treated | 29 | Male | Asian | ABC/TCV/3TC | <40 | 547 | 444 |
| 8011* | Acutely-treated | 46 | Male | Hispanic/Latino | BIC/FTC/TAF | <40 | 738 | 386 |
| 8012 | Acutely-treated | 49 | Male | White | TCV, TRU | <40 | 478 | 471 |
| 8019* | Acutely-treated | 31 | Male | White | RPV/TAF/FTC, TCV | <40 | 448 | 354 |
| 8039 | Acutely-treated | 24 | Male | Mixed Race/Multiracial | ABC/TCV/3TC | <40 | 405 | 598 |
| 8053* | Acutely-treated | 35 | Male | Mixed Race/Multiracial | BIC/FTC/TAF | <40 | 516 | 1059 |
| 8083 | Acutely-treated | 27 | Male | Mixed Race/Multiracial | BIC/FTC/TAF | 108 | 166 | 319 |
| 1379* | Chronically-treated | 50 | Male | Black/African American | DRV, RTV, TRU | <40 | 522 | 596 |
| 2021 | Chronically-treated | 52 | Male | Black/African American | RPV/TAF/FTC | <40 | 500 | 857 |
| 2161* | Chronically-treated | 71 | Male | White | DRV, RTV, TCV, 3TC | <40 | 568 | 479 |
| 2253 | Chronically-treated | 69 | Male | White | BIC/FTC/TAF | <40 | 349 | 354 |
| 2553* | Chronically-treated | 36 | Female | Hispanic/Latino | ABC/TCV/3TC | <40 | 562 | 721 |
| 3037* | Chronically-treated | 61 | Male | Black/African American | BIC/FTC/TAF, DRV, DOR | 54 | 598 | 1326 |

**Abbreviations:** BIC: Bictegravir; FTC: Emtricitabine; TAF: Tenofovir alafenamide; ABC: abacavir; TCV: Tivicay; 3TC: lamivudine; TRU: Truvada; RPV: Rilpivirine; DRV: Darunavir; RTV: Ritonavir; DOR: Doravirine. \*Indicates those participants from which paired blood and lymph node specimens were collected.

**Table S2.**
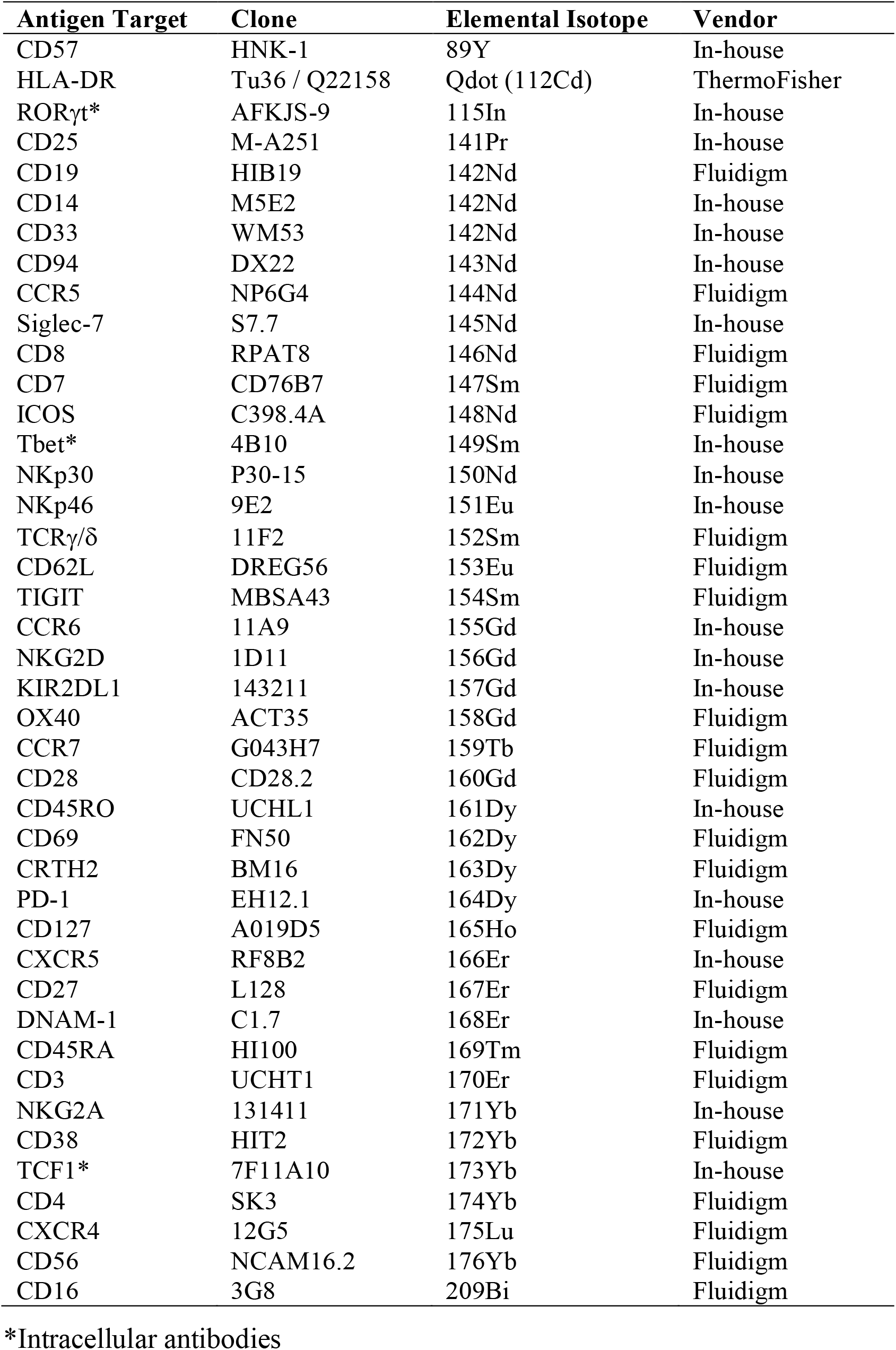
List of CyTOF antibodies used in this study. Antibodies were either purchased from the indicated vendor, or prepared in-house using commercially available MaxPAR conjugation kits according to the manufacturer’s instructions (Fluidigm).

